# Comparing the predictors of mutability among healthy human tissues inferred from mutations in single cell genome data

**DOI:** 10.1101/2023.11.28.569048

**Authors:** Madeleine Oman, Rob W. Ness

## Abstract

Studying mutation in healthy somatic tissues is key for understanding the genesis of cancer and other genetic diseases. Mutation rate varies from site to site in the human genome by up to 100-fold and is influenced by numerous epigenetic and genetic factors including GC content, trinucleotide sequence context, and DNAse accessibility. These factors influence mutation at both local and regional scales and are often interrelated with one another, meaning that predicting mutability or uncovering its drivers requires modelling multiple factors and scales simultaneously. Historically, most investigations have focused either on analyzing the local sequence scale through triplet signatures or on examining the impact of epigenetic processes at larger scales, but not both concurrently. Additionally, sequencing technology limitations have restricted analyses of healthy mutations to coding regions (RNA-seq) or to those that have been influenced by selection (e.g. bulk samples from cancer tissue). Here we leverage single cell mutations and present a comprehensive analysis of epigenetic and genetic factors at multiple scales in the germline and three healthy somatic tissues. We create models that predict mutability with on average 2% error, and find up to 63-fold variation among sites within the same tissue. We observe varying degrees of similarity between tissues: the mutability of genomic positions was 93.4% similar between liver and germline tissues, but sites in germline and skin were only 85.9% similar. We observe both universal and tissue-specific mutagenic processes in healthy tissues, with implications for understanding the maintenance of germline versus soma and the mechanisms underlying early tumorigenesis.

**Summary:** Mutations in healthy tissues can reveal how genetic diseases originate. In this study, we explore how mutation rates vary across the human genome and what influences these variations. We leverage advanced single-cell analysis to analyze genetic and epigenetic factors in germline and three healthy tissues. We trained models that exhibit high accuracy and discover large variation within the same tissue. We also identified differences in mutability between tissues, suggesting both universal and tissue-specific mutation patterns with implications for understanding oncogenesis.

## Introduction

Mutation is a continual process that occurs in both somatic and germline tissues. The consequences of germline mutation are widely appreciated as they underpin both heritable genetic disease and adaptive evolution. The consequences of somatic mutation are similarly profound including the origin of cancers (García-Nieto et al. 2019), neurodegeneration (Lodato et al. 2015) and ageing (Finkel et al. 2007). Although mutation is a random process, the likelihood of mutation (mutability) at any given site is not uniform across the genome. Variation can be seen in large genomic windows but also can vary between adjacent sites, and the rate at which mutations occur can vary from site to site more than 100-fold (Ellegren et al. 2003).

Mutability variation is driven by both genetic and epigenetic processes. A well known example of a genetic effect on mutability is CpG dinucleotides which are 10-fold more likely to mutate than non CpG sites (Hwang and Green 2004; Hodgkinson and Eyre-Walker 2011; Oman et al. 2022), when methylated cytosines are converted to thymine through deamination (Coulondre et al. 1978; Razin and Riggs 1980). Numerous epigenetic processes have also been associated with mutability variation. For instance, regions close to the origin of replication are associated with lower mutation rates, due to the increased activity of DNA repair (Supek and Lehner 2015). Similarly, transcription level, chromatin conformation, various histone marks, and recombination (Michaelson et al. 2012; Schuster-Böckler and Lehner 2012; Blokzijl et al. 2016; Carlson et al. 2018; García-Nieto et al. 2019; Moore et al. 2021) have also been linked to mutation rate variation. It is important to note that many of these genetic and epigenetic processes that affect mutability are interdependent (Roadmap Epigenomics Consortium et al. 2015), so if we hope to determine the relative contribution of these different processes on mutation rate variation we need to consider them together in models that can account for correlations among these forces (Chen et al. 2017; Supek and Lehner 2019). For example, though DNAse accessibility correlates negatively with mutation at the megabase scale (Polak et al. 2013), this effect was found to be minimal after controlling for other confounds (Supek and Lehner 2017). Specifically, DNAse accessibility was identified as a predictor for replication timing (Gindin et al. 2014). Studies that have analysed multiple epigenetic and genetic factors simultaneously (Chen et al. 2017; García-Nieto et al. 2019) have analyzed larger genomic scales such as on individual genes. Consequently, these studies do not capture the significant variations observed at smaller scales, including per base variation. This limitation is critical given that small-scale sequence context has been shown to explain 81% of the variance in human substitution probabilities (Aggarwala and Voight 2016), emphasizing the necessity for analyses that include finer resolutions. The few studies that incorporate multiple scales have only modelled the human germline (Michaelson et al. 2012; Seplyarskiy et al. 2023), whereas these analyses need to extend to somatic tissues to understand the predictors of mutability in all healthy tissues types across the human body.

Cellular environments and epigenetic landscapes impose substantially different mutagenic pressures across tissues and cell types (Roadmap Epigenomics Consortium et al. 2015; García-Nieto et al. 2019). This is in evidence from a large variation in mutation load among tissues, ranging 23-fold from the highest in intestinal crypts to the lowest in germline (Ren et al. 2022). However, the forces responsible for this variation and its full extent across the body remains unclear. The mutational landscape of somatic tissues is an active area of study, however the literature on mutation rate variation in healthy tissues is relatively sparse. Studies focusing on healthy tissues are limited by the inherent difficulty of sequencing rare, mosaic somatic mutations. Until recently, our knowledge of somatic mutation has relied primarily on bulk sequencing of tumours (Schuster-Böckler and Lehner 2012; Alexandrov et al. 2013), which can only extract mutations that occur at relatively high frequency and therefore have been influenced by clonal selection (Malikic et al. 2019) and may be biassed due to changes in the cellular and mutagenic environment when tissue becomes cancerous. Other studies have used RNA (García-Nieto et al. 2019) or exome sequencing (Chen et al. 2017), and thus can detect patterns present in the coding regions but can not analyse the majority of genome space that is non-coding. Single cell sequencing (SCS) is a technology that may offer a method to identify low frequency mutations in healthy tissue. SCS has the power to greatly increase the quantity and variety of discoverable mutations, facilitating the analysis of the complete spectrum of somatic mutations and providing an unbiased view of the mutations and their drivers across tissues. Until recently SCS has been prohibitively costly (Wang and Navin 2015), and there has therefore been no studies that use SCS mutations to analyse multiple epigenetic and genetic predictors of mutation at fine resolution across germline and soma tissues (Supek and Lehner 2019).

To address this gap, we built multivariate regression models for four healthy tissues that predict the relative probability of mutation at individual sites from 21 epigenetic and genetic predictors. We leveraged pre existing datasets, including high quality SCS datasets for blood, liver and skin tissues, and parent-offspring trios mutations in germline. We explored how somatic mutational processes vary across healthy human tissue, and specifically addressed the questions:

1. How accurately can genetic and epigenetic processes predict mutation rate variation in different healthy tissues?
2. What genetic and epigenetic factors predict mutation in healthy tissue?
3. How do the factors that predict mutation rate vary between tissues?
4. How much does the mutability of a site vary among tissues?

## Results

### How accurately can genetic and epigenetic processes predict mutation rate variation in different healthy tissues?

Our mutability models accurately predicted mutability in all four tissues analysed. In blood, germline, liver and skin we report low error (average Mean Absolute Error, MAE = 0.0215) based on the fit of predicted mutability with the proportion of mutations in each bin of sites (figure 1). The fact that mutability learned from a training set was able to predict the proportion of mutations in each bin of independent test datasets demonstrated that the genetic and epigenetic properties can predict the main mutagenic processes occurring in blood, liver, germline and skin. We see a wide spread of predicted mutability, ranging from 0.2 to 0.9, implying a wide range of actual mutability among sites. The wide range of mutability is another indicator that the model is making meaningful predictions because in the absence of a strong signal the null estimate for each site would be 0.5 mutability since our dataset contained 50% mutations. Extreme mutability values that deviate from this null demonstrate that the model is able to differentiate high and low mutability sites; indeed, when we randomize the mutation status of the training data, mutability values cluster strongly around 0.5 and 73% of sites have mutability between 0.45 and 0.55, in contrast to only 18% of sites in this range in the real data (see figure S1).

**Figure 1:**
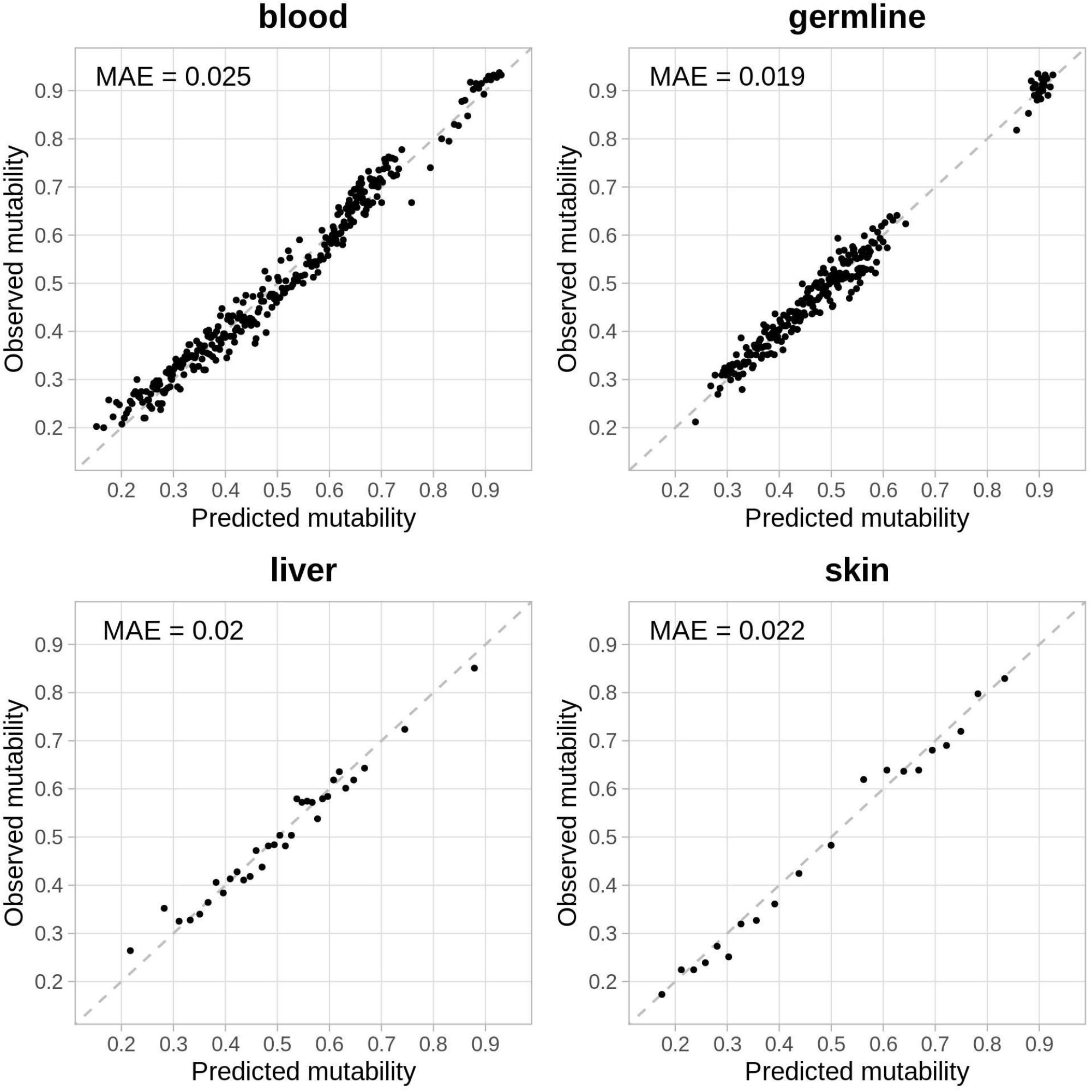
Tissue specific mutability model fit and accuracy in healthy tissues. Here accuracy is estimated from how well the estimated mutability predicts the proportion of the sites in a given bin that are actually mutations as mean absolute error (MAE). Each data point represents a bin of 400 mutated or unmutated sites within a given mutability range. The proportion of sites that are true mutations in that bin is plotted against the average predicted mutability, which represents the model’s estimation of the probability that a site would be a mutation in the test set. The dashed grey line is the 1:1 where the predicted mutability and proportion of mutations are equal.

Interestingly, in the germline model we see a considerable gap between the highest mutability sites and the rest, which appears to be driven by the extreme difference in mutability of germline CpG sites. CpG sites are known to be extremely mutable and so to test if these sites alone are driving model accuracy across tissues, we created models removing these sites. Without CpGs, the spread of germline mutability predictions reduces 40% from 0.24-0.93 to 0.27-0.67, however, the model maintains the same low error on the remaining sites (MAE=0.02). MAE remains low in blood, liver and skin when CpGs are removed (mean MAE = 0.022). No other tissue exhibits the same reduction in the spread of mutability values as in the germline, with the skin only experiencing a 2.5% decrease from 0.17-0.97 to 0.17-0.95 (see figure S2). Thus we find that, while CpG conversion accounts for the main mutagenic process in highly mutability sites in all tissues but skin, there are numerous other critical mutagenic processes that we have identified to accurately predict the ratio of mutant and non-mutant sites at lower mutability.

### What genetic and epigenetic factors predict mutation in healthy tissue?

The impact of each epigenetic and genetic factor on mutability can be extracted from each tissue model by analysing t-statistics. The t-statistic is a standardised parameter that indicates the strength and direction of effect a given predictor has on mutability, controlling for the confidence in that value. Across tissues, sequence triplets consistently emerge as strong predictors of mutation, with both mutagenic and protective effects seen for different triplets (figure 2). Uniformly across tissues we find that CpG sites (XCG triplets) are positively associated with mutation, however this effect is strongest in the germline, where CpG have two fold higher mutability than non CpG triplets. Furthermore, t-statistics for CpG triplets are 20% higher in the germline than other tissues. Another significant pattern is that all triplets with a central T (or A in the complementary strand) are significant and negatively associated with mutation (denoted here as “XTX”). This effect varies among XTX triplets, with some triplets exhibiting strongly protective effects against mutability (e.g. mean t-statistic for TTT across tissues= -7.7±2.5), while other triplets have an effect close to zero (e.g. mean t-statistic for ATC across tissues = -1.6±2.1). This trend is seen in all tissues, however the protective effect of XTX triplets is much stronger in the blood: t-statistics for XTX triplets are 3.2 times more negative in blood compared to germline, skin and liver. A third triplet signature we find is that [T/C]CX triplets are extremely strong positive predictors of mutation in skin (4.6 times higher t-statistic values than all other triplets), but this effect is not seen in other tissues, likely due to the fact that mutations at these sites are typical of UV light damage.

**Figure 2:**
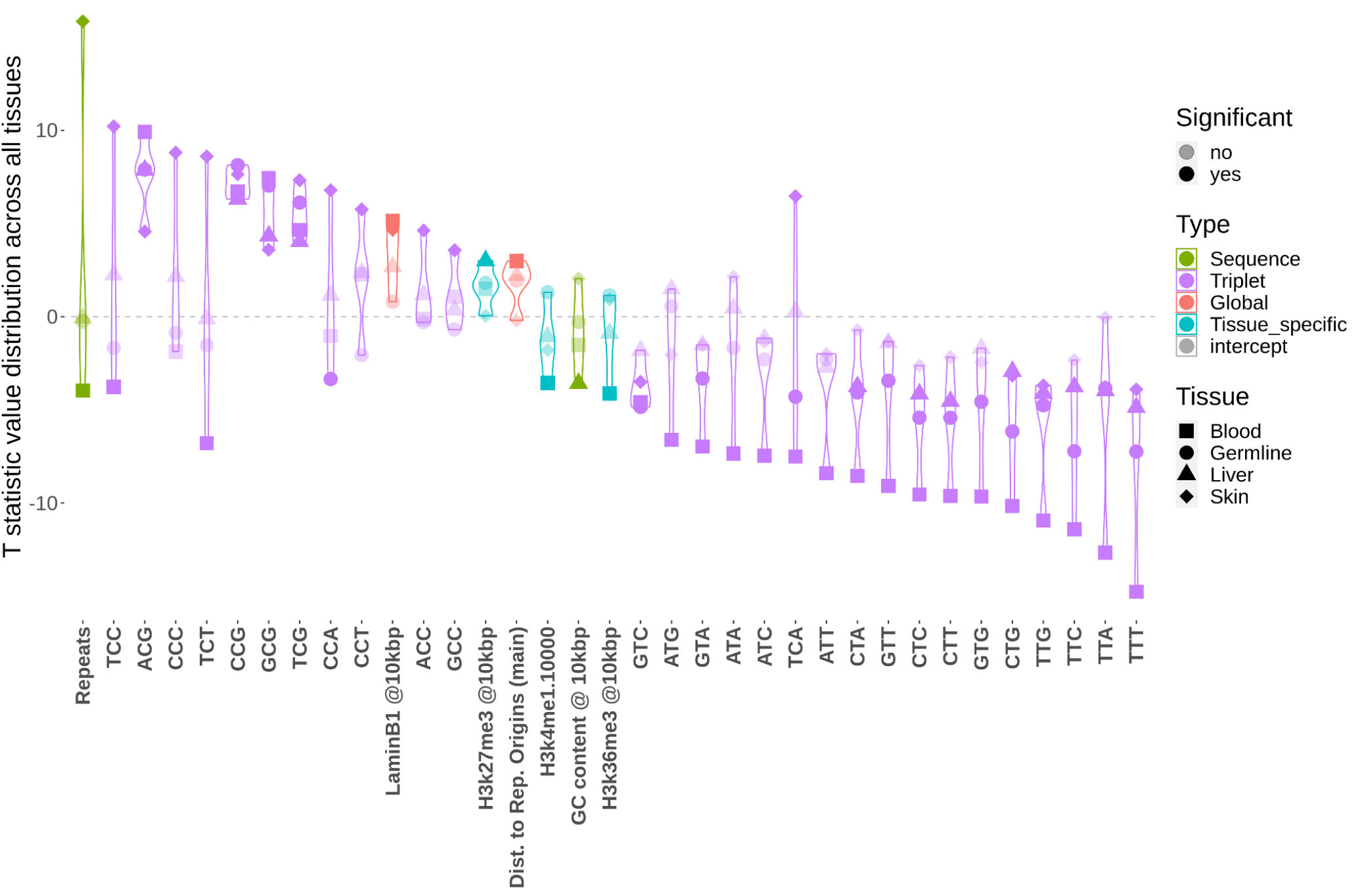
Comparison of the importance of genomic and epigenetic predictors of mutability among tissues. Every predictor is represented by four points corresponding to each of the tissues, where each point represents the t-statistic for a given tissue. The larger deviations from zero (grey dotted line) indicate a stronger effect on mutability, where points above the line increase mutability while those below reduce mutability. Here we only plot predictors that have a significant effect (p <0.005) in at least one tissue; tissues in which a given genomic property has no significant effect are indicated with translucent data points. Predictors are ordered by the the most extreme t-statistic value of the four tissues from most positive to most negative. A predictor with the same mutagenic importance in two tissues would exhibit a larger t-statistic in the tissue with the larger dataset. Thus, to ensure discrepancies in t-statistics reflect biologically meaningful differences, here the germline, liver and blood datasets are downsampled to be equivalent to the smallest dataset, skin (n mutations = 10,191). See figure S8 and table S9 for t-statistics values from the full models without downsampling.

We see that a diversity of epigenetic and genetic factors are important for predicting mutation beyond triplets. Being in repetitive DNA is strongly indicative of mutation in the skin (t-statistic = 15.9). Interestingly, while the effect of repeats in skin is the most mutagenic effect seen across all tissue models, repeats have close to no effect on the liver or germline mutability (t-statistics = -0.07 and -0.26 respectively), and is protective in the blood (t-statistic = 4.0). The GC content of the surrounding 10Kbp protects against mutation, with a strong negative influence on the predicted mutability of a central site. However, this trend is only seen in the liver (t-statistic = -3.6), with no significant effect in blood, skin or germline (t-statistic = -1.53, 2.04 and -0.28 respectively) . Epigenetic processes are also important to predicting mutation. Here we find that sites at the nuclear periphery (indicated by high laminB1 values) are more likely to mutate, though this effect is only significant in the blood and skin tissue (t-statistic = 5.13 and 4.66 respectively). Histones marks are also important, though typically only significant in one tissue; H3k27me3 marks (indicating facultatively repressed genes) are mutagenic in the liver (t-statistic = 3.0), while H3k36me3 marks (actively transcribed genes) are protective in the blood (t-statistic = -4.12). The variation seen in the importance and direction of effect of epigenetic and genetic factors included in the model suggests that mutagenic processes are specific to the cellular environments within each tissue.

Triplets play an important role in the predictive ability of these models, however we wanted to test whether model accuracy relies solely on these predictors. Thus we created models without triplet predictors and evaluated accuracy to see if other genetic and epigenetic factors contain enough biologically relevant signal to accurately predict mutability. Models without triplet predictors still exhibit high accuracy, with MAE only increasing on average 12% from 0.0215 to 0.024 across tissues. Recall that another indicator of how well the mutability model is working is the range of mutability values predicted: accurate extreme predictions (close to either 1 or 0) indicate that the model has the power to leverage the epigenetic and genetic factors to assign a site high or low mutability, while models that do not find associations between the predictors and mutation will default to predicting 0.5 mutability as there are 50% mutations in the dataset. Without these predictors, we find that the spread of predicted mutability drops 42% across tissues. In the absence of triplets, we find the emergence of other predictors that can capture mutagenic signals (see figure S4). Across tissues we find that GC-content at the 100 scale is mutagenic (average t-statistic = 7.7±3.0), possibly reflecting some of the signal from high mutability CpG triplets. Similarly, sites at the nuclear periphery are universally mutagenic in the absence of triplets (average t-statistic= 3.8±0.6), as opposed to the tissue specific effect seen in the full model (see figure 2 above). Models without triplets also reveal robust tissue-specific signals: for example, repetitive sequences exhibit pronounced mutagenicity in skin (t-statistic = 13.7), yet confer a protective effect in the germline and blood (t-statistic = -5.9 and -14.0 respectively). Similarly, many histone marks become important for predicting mutation but only in one or two tissues; for example sites enriched for H3k27me3 is strongly mutagenic in the germline and liver (t-statistic = 8.3 and 5.9 respectively), but has close to no effect in the blood or skin (t-statistic = -2.4 and -0.02 respectively). The fact that models without triplets maintain accuracy demonstrates that, while triplets are important indicators of mutagenic processes, these mutagenic signals are also encompassed within diverse genetic and epigenetic factors.

Notably, we included many genomic properties and genetic processes in our models that have been found to drive mutation in other studies, but found no significant predictive power for mutability here. For example, recombination rate has been shown to correlate with mutation (Lercher and Hurst 2002) due to imperfect repair at recombination sites. However here we find no importance of recombination in the germline tissue, with a t-statistic of 2.07 and p value of 0.038; though traditionally this would be considered significant, here we consider over 300 predictors across all tissue models and so stricter significance cutoffs are required to protect against type 1 error. Similarly, distance to replication origin has been correlated to human SNP density (Wolfe et al. 1989; Stamatoyannopoulos et al. 2009; Chen et al. 2010) and mutations in cancer genomes (Woo and Li 2012; Liu et al. 2013; Supek and Lehner 2017), in part due to the fact that DNA repair machinery is recruited to regions that replicate early (Supek and Lehner 2015). Again we find no significant effect of replication timing in mutation in liver, germine or skin models (p values range from 0.027 to 0.208).

### How do the factors that predict mutation rate vary between tissues?

We can analyse directly how the mutagenic effects of epigenetic and genetic factors vary between tissues by comparing the t-statistic between models in a pairwise fashion: if predictors have similar mutagenic effects in the two tissues being compared, then the data will be near the 1:1 line with little dispersion. We find that the process of mutation in skin is distinct from other tissues. There was a lower correspondence of t-statistics, with an average MAE of 0.096 compared to the other three tissues, which is a 1.7x larger difference than observed between other tissue pairs (see table 1). We can see the extent to which mutability differs in skin in figure 4B where many genomic and epigenetic properties that are negatively associated with mutation in blood, are positively associated with mutation in skin (MAE=0.091). In contrast, the greater similarity in the strength and direction of mutagenic processes in other tissues is exemplified in figure 4A. When comparing the germline and liver, we find a linear relationship between the t-statistics, though they do not always lie along the 1:1 line. We see low error (MAE=0.044) with only GC content switching in its direction of effect on mutation between the tissues. Interestingly, the pairwise comparison reveals that predictors in the blood are generally lower than in other tissues (t-statistics are 3.5 times lower than in other tissues, ±1.2). Comparisons between all tissues can be found in the supplemental figures S6-7 and while they show varying degrees of similarity between the blood, liver and germline, it is clear that the skin tissue is the most unique among those compared here.

**Table 1:** Summary of the overall similarity of t-statistic values between tissues. Below we report the mean absolute difference of the corresponding t-statistic for a given pair of mutability models, corresponding to figures 4, S6 and S7.. Larger numbers indicate bigger discrepancies between the mutagenic effect of different genetic and epigenetic factors across tissues.

|  | germline | liver | skin |
| --- | --- | --- | --- |
| blood | 0.053 | 0.065 | 0.114 |
| germline |  | 0.051 | 0.094 |
| liver |  |  | 0.079 |

**Figure 4:**
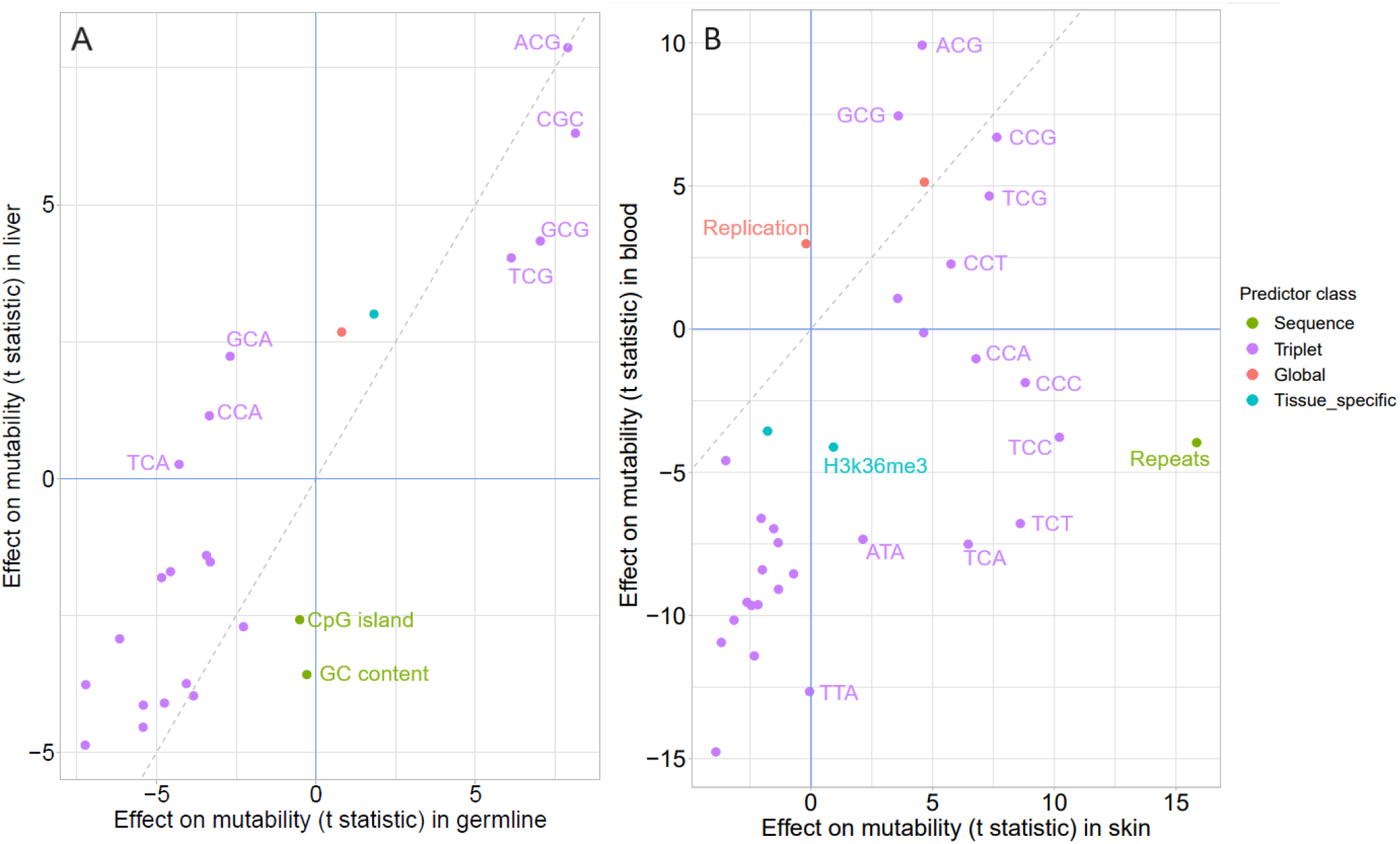
Pairwise comparisons of the strength and direction of different genetic and epigenetic properties and processes on mutability across tissues. Only predictors that are significant in at least one of the tissues after correcting for multiple tests are displayed (p<0.01). These pairwise comparisons were chosen to illustrate how the trends differ between similar models (A. germline vs. liver) and markedly different models (B. blood vs. skin). Predictors that have a similar effect in both tissues are near 1:1 line (grey dotted line), while those that deviate from equality have an effect unique to that tissue. Predictors in the top left or bottom right quadrant change the direction of effect between tissues, indicating a process that is mutagenic in one tissue and protective in another. Select informative points are labelled. Because the magnitude of t-statistics are proportional to the size of the data set we downsampled each tissue’s training set to be equivalent to the smallest dataset for this analysis (skin n_mutations_ = 10,191). For comparison of other tissue pairs, see figures S6-S7.

To test if tissue specific patterns arise as artifacts of different experimental methods used between tissues, we tested the blood tissue model on an independently generated dataset of single cell blood mutations (n=9,181, full details in table S1). We find a low error rate (MAE=0.035), supporting the validity of our method (figure S9). The low error aligns with the similarity observed when we repeat this comparison with the blood and liver models (MAE=0.031), indicating that the mutability between these tissues is analogous to that within a single tissue sample. However, when examining other tissues, we noted a more than twofold increase in error (average MAE=0.075), indicative of distinct mutagenic signatures specific to each tissue type (see figure S5).

### How much does the mutability of a site vary among tissues?

We compared the predicted mutability of the same 15000 random sites estimated from each of the tissue specific models to determine the consistency of mutation rates across the genome in different cellular contexts. Overall the mutability of sites was clearly correlated across tissues with a mean MAE of 0.10 (figure 5) but the errors varied from 0.141 between skin and germline mutability to 0.064 in germline versus liver. Differences in the strength of correlation were driven by strong patterns of clustering, especially in skin and germline tissues. In all tissues CpG sites are enriched for high mutability, but the effect is exaggerated in germline where the mutability of CpG sites is separated from the rest of sites (figure 5, purple). In skin the [T/C]CX triplets, likely the result of UV damage, are distinct in that they have high mutability in skin and relatively average mutability in other tissues (figure 5, blue). There are also a group of sites from repetitive DNA that have very high mutability in skin (mutability >0.9), but are not very mutable in the other tissues. Therefore, while there are correlations in the mutation rate across tissues, strong patterns are apparent that suggest the same site may have very different mutation rates depending on the cell it finds itself in.

**Figure 5:**
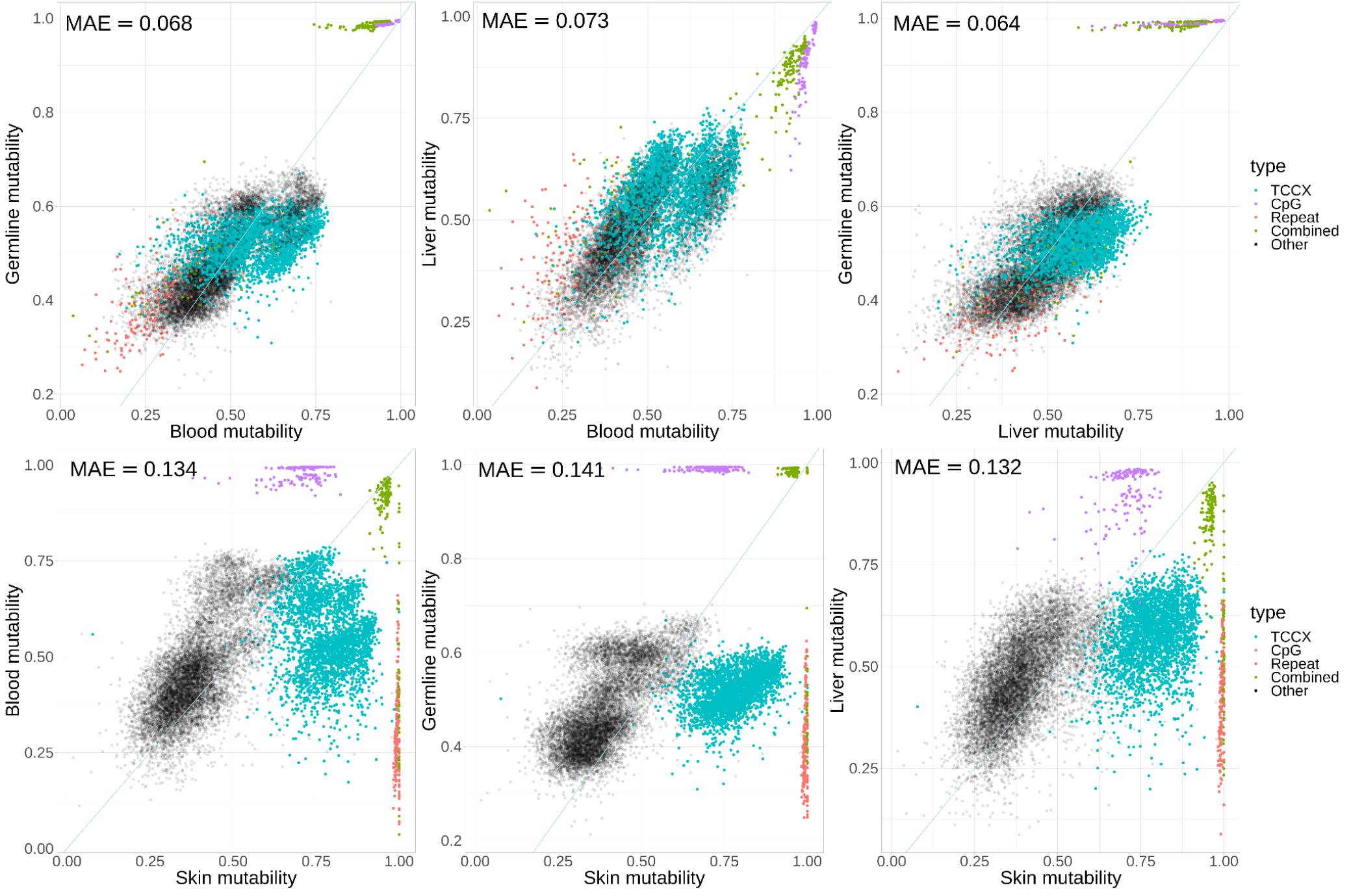
Pairwise comparisons of per site mutability in different tissues. Blood, germline, liver and skin models were all used to predict mutability on a set of 15000 random sites. Sites are coloured to highlight the key factors that result in mutability variation between tissues, including CpG triplets, [T/C]CX triplets, and DNA repeat context. Any site that was a combination of these categories was labelled ‘combined’ and all other sites are shown as translucent black points. The 1:1 line is shown in grey and the mean absolute error (MAE) from that line is reported in each panel.

## Discussion

### Mutability models predict 63-fold variation in mutation rate with high accuracy

Our goal was to leverage single cell mutations to explore the effects of genetic and epigenetic factors together on mutability at the single base resolution in healthy human tissues. We compiled the main genomic factors known to affect mutation and created mutability models for the germline and soma (blood, liver and skin) that all demonstrate impressive precision and low error (average MAE = 0.0215). In all tissues we successfully discriminated between highly mutable and less mutable sites. For example, 90% of sites classified to have a mutability score of 0.9 are indeed true mutations, even though the accuracy was tested on an independent test set and only 50% of test sites were mutations. The converse is also true, as these models can successfully classify non-mutant sites into lower mutability bins where there are very few true mutations (as low as 17.3%). Note that non-mutant sites in high mutability bins and mutant sites in low mutability bins are not necessarily misclassified; instead it is expected that most high mutability sites in the genome do not mutate regularly and some low mutability sites do mutate despite this being relatively less probable. This fact is what necessitates binning sites by predicted mutability to evaluate the performance of the models. The fact that predicted mutability correlates strongly with the observed fraction of mutations suggests the mean mutability of sites in each bin is well estimated, even with narrow ranges of mutability in bins and relatively few sites (400 sites). This does not negate the possibility of some inaccuracy and unexplained variance in the predictions of mutation rates of individual sites (Eyre-Walker and Eyre-Walker 2014). Very similar approaches have been used to predict mutability in model organisms (Ness et al. 2015; Saxena et al. 2019; López-Cortegano et al. 2021) and the human germline (Michaelson et al. 2012; Seplyarskiy et al. 2023) but our work represents the first time this approach has been used in human soma. The accuracy of these models reinforces the utility of this statistical approach, and that mutability can be predicted from models trained with as few as 10,191 mutations. Importantly, the fit of the models demonstrates that the epigenetic and genetic predictors included here capture important mutagenic signals.

### DNA sequence-based properties are the strongest predictors of mutation rate variation across all tissues

We find that many previously known trends hold true in healthy tissue when leveraging unbiased SCS mutations. In our analysis, the DNA sequence itself emerges as the strongest predictor of mutation. Historically, studies have analysed the impact of genetic and epigenetic changes on mutation rate variation in large genomic regions (Chen et al. 2017; García-Nieto et al. 2019). In contrast we find that the local scale (<10bp) is the most important for predicting mutation. In line with previous observations, both the specific nucleotide at a site and its two flanking sites (triplet) strongly predict human mutability (Michaelson et al. 2012; Aggarwala and Voight 2016). Among triplets, we find a wide range of mutagenic effects, with the t statistics of the most extreme triplets (ACG=29.4, TTT=-49.2) over 3.5 standard deviations away from the average (mean=-5.8, sd=13.9). As expected, deamination of methylated cytosines causes CpG sites to be highly and universally mutable, however the strength of this effect across the four different CpG triplets (XCG) varies up to 2.4 fold within a tissue. Conversely, we find that triplets with a central A/T (denoted X[A/T]X) are protective against mutation across all tissues, but with over 7-fold variation between the most and least protective X[A/T]X triplet within a tissue. Our findings suggest that local sequence context is one of, if not the best predictor of mutability in healthy tissue and can also drive changes in the mutation rate of the same site in different cell types. This argues that models of sequence evolution should be using the sequence context to determine the null expectation of mutation rate variation, but each species and cell type will require its own dataset to completely capture the variation in mutation rate in different contexts.

One of the reasons that triplets are such a powerful predictor of mutation is because they capture many different mutagenic processes that affect DNA in a biassed way (Alexandrov et al. 2013; Moore et al. 2021). These effects can be due to different repair mechanisms or variable agents that cause specific types of DNA damage. This means that although the sequence is the same across tissues, the triplets can have different mutation rates depending on the specific cellular environment. For example, the protective effect of triplets with an A/T at the central base is 3.2x stronger in blood than all other tissues, suggesting that transient strand denaturation at A/T sites may play a lesser role in mutagenesis in blood (Leroy et al. 1988). We also find that CpG sites are 20% more mutable in the germline than soma (see below). Variable triplet mutation rates have been shown to reflect numerous external mutagens (Alexandrov et al. 2013). A clear example is UV light that causes mutations in triplets with a central C and either a C or T upstream (denoted [T/C]CX). Consequently, we find [T/C]CX triplets are highly mutable in skin, while having a protective or inconsequential effect in other tissues. Interestingly, this effect is seen in the skin regardless of chromatin accessibility, suggesting that mutations from UV damage accumulate independent of condensation state (Pich et al. 2018). Thus, consistent with other work (Alexandrov et al. 2013; Moore et al. 2021), we find that triplets are important for capturing the mutagenic processes that are endogenous to cells, as well as those incurred via external mutagens.

### Epigenetic factors that capture mutagenic processes vary among tissues

Though triplets are powerful predictors of mutation, other genetic and epigenetic factors also account for a significant fraction of the mutagenic signal. We created models without triplets and found that models lose the ability to predict the extremes of the mutability distribution, with a mean 43% reduction in the range of predicted mutability (see table S3). However, models still retain accuracy, as MAE increases only 7.5% within this range. The importance of epigenetics factors is in line with other studies that find that such factors can explain between 55-80% of mutation variation in cancer (Schuster-Böckler and Lehner 2012; Polak et al. 2015). Many internal processes of different cells are interdependent and involve interactions of DNA with epigenetic mechanisms, thus both the sequence-based and epigenetic predictors may capture the same mutagenic processes. When all predictors are combined triplets emerge as superior predictors, likely owing to the fact they can capture the notable local-scale variability in human mutation rates. However, the variation in triplet mutability among the limited set of tissues analyzed here argues for much more comprehensive sampling of different cell types, individuals, ages and even species if we hope to produce accurate predictive models of mutation.

Despite this, the effect of epigenetic processes on mutation provides both insight into the mechanism of mutation and captures information not reflected by triplet sequence. In general, we find that open chromatin regions are associated with lower mutability. Regions at the nuclear periphery (laminB1) are silenced as heterochromatin (Filion et al. 2010), and being in these regions is a strong indicator of mutation in blood and skin. In both germline and liver tissue, sites in facultatively repressed chromatin (H3k27me3) are associated with higher mutability (t-statistics =5.4 and 5.7 respectively) while in the blood, regions that are annotated (t-statistic = -2.8) and marked (H3k36me3, t-statistic = -4.0) as genes have lower mutability. Our models therefore support the hypothesis that transcription coupled repair outweighs the damage caused by transcription (Moore et al. 2021). However, the fact that we do not see as strong and consistent a signal across tissues suggests that the relationship between chromatin state and mutability may be more nuanced than previously thought or not particularly strong. In fact, other previously established trends are weaker or have variable effects across tissues in our analyses. For example, we only see an effect of replication timing and H3k36me3 in blood, whereas these predictors have previously been strongly linked with mutability in cancer (Woo and Li 2012; Liu et al. 2013) and in RNA-seq data (García-Nieto et al. 2019). These discrepancies highlight the importance of studying the whole genome, and how using single cell mutation data is critical to revealing the mutagnic processes at work in the early stages of genetic disease. The fact that the same predictors have variable effects across tissues suggests that there are important tissue specific effects that influence mutability and contribute to the overall mutation rate and spectra in each tissue.

### Mutability in healthy tissues exhibit varying degrees of similarity

We also explored the similarity of mutability across healthy human tissues by comparing how the effect of each epigenetic and genetic predictor, how similar model predictions were between tissues and how this leads to variation in mutability of sites across tissues. The first major finding is that the mutation profile of skin is the most distinct, driven primarily by the preponderance of mutations with the triplet signature of UV damage, exemplifying how external mutagens can dominate mutation. Interestingly, the liver and blood models can predict mutations in skin with only moderately reduced accuracy. This suggests that, besides the external mutagen UV light signature, the mutagenic processes that occur in these tissues are similar, and that the liver and blood models are accurately predicting mutations not caused by UV damage in skin. This is supported by work that extracted the triplet signatures in healthy tissue and found that the two endogenous triplet signatures that dominate many tissues are also present in skin (Moore et al. 2021). The similarity between skin and liver may be further explained by oxidative damage in the liver, as [T/C]CX triplets are often mutated in the signature associated with APOBEC (Alexandrov et al. 2013).

The second major finding is that the blood and liver tissues have similar mutability while the germline tissue is distinct, though this difference is less extreme than that seen in skin. The blood tissue model exhibits the same degree of accuracy when predicting on liver as an independent blood dataset, while the germline model exhibits only moderate accuracy (figure S9). This suggests that mutability in the liver and blood tissues is just as similar as between two samples of the same tissue. The feature likely driving the difference to germline is the relatively elevated mutability seen in CpG sites: we find the effect is three times greater than in other tissues. Interestingly, in an earlier study investigating the effect of triplet sequence on mutation they found a similar contribution of the CpG signature in germline and soma (Moore et al. 2021), and report in general that the major triplet signatures were shared across most tissues. In the two studies that have examined the effects of epigenetic processes on mutation in human germline and soma, differences between the soma and germline were identified (Chen et al. 2017; Moore et al. 2021). Specifically, transcription played a protective role in soma (over ten tissues including skin and liver) but not in germline, and the correlation between late replicative timing and increased mutation was stronger in soma. Notably these associations were based on mutations in coding sequence only that were modelled with one variable at a time (Moore et al. 2021) or from mutations from bulk cancer tumours (Chen et al. 2017), and these differences could explain why we did not observe either pattern in our data. The fact that our results differ highlights the importance of including local scales, analysing intergenic regions, and modelling multiple variables simultaneously to control for correlations among genomic properties. These differences between germline and soma are worth noting because analyzing patterns of genetic variation in a pedigree, a population or through evolutionary time reflects the process of mutation in the germline, and those findings may bias our assumptions about mutability across the body and how it influences somatic mutation and disease.

Differences in mutation profiles across various cancers have been well established (Alexandrov et al. 2013; Chen et al. 2017), but the time when these changes arise remains unclear. It raises a critical question: to what extent do mutation differences among tumours reflect mutability in healthy tissues versus changes in mutability acquired during tumour evolution? Comprehensive analyses of healthy somatic tissues are scarce. (García-Nieto et al. 2019) detected differences between tissues using epigenetic and genetic factors, however, their analysis was limited to RNA-seq data, capturing only the 1.5% (Lander 2011) of the genome in coding regions. (Moore et al. 2021) found that genome-wide, two triplet signatures accounted for the majority of mutations in most tissues, arguing that healthy tissues generally share mutation processes. However, in a limited analysis of the impact of gene expression and replication timing, Moore et al. did find differences between germline and soma tissues. Our work represents the first time mutations from single cell sequencing have been leveraged to analyse multiple epigenetic and genetic predictors of mutation at fine resolution across germline and soma tissues (Supek and Lehner 2019). By collating data from four tissues and twenty one genomic properties we have shown that somatic mutability is both complex and differs among tissues. Our comparison of mutability of the same sites in different cellular contexts (figure 5) clearly demonstrates that sites mutate at very different rates in different cells. Therefore, the probability of a site contributing to ageing and disease across tissues is highly variable. If this variation results in some genes being more prone to mutation, it could mean that their association with disease could be driven by higher mutation rate, by causing the disease, or both. We are currently limited by somatic sequencing technology: many tissues are not amenable to the current standard practices, because they cannot be clonally expanded (e.g. post mitotic tissues like the brain) or because they do not have local monoclonal structures available for for laser-microdissection (e.g. the blood). Error corrected sequencing technologies have reported error rates as low as 5 in a billion bases (Abascal et al. 2021) and will offer a new way to identify somatic mutations. Analysing larger datasets and more healthy tissues than ever before will help deconvolute the mutagenic processes that occur at the earliest stages of disease and can identify targets for preventive treatment.

## Methods

### Mutation data sets

To do a comprehensive analysis of how epigenetic and genomic processes impact mutability in four healthy tissues we leveraged publicly available mutation datasets from pedigree and single-cell sequencing. We used 104,941 single nucleotide mutations in the germline, identified from whole genome sequencing of 1.5k parent-offspring trios (Jónsson et al. 2017). For somatic tissues, mutations from SCS were downloaded from the Database for Somatic Mutations in Normal Cells (Miao et al. 2019) or the database of somatic mutations in normal tissues (Sun et al. 2022), see supplemental table 1 for a summary. Mutation datasets varied in size (blood = 9,082, germline=104,941), however in each model we included the same number of non-mutant sites as mutant, assuring 50% mutation frequency across all tissues (see the section Training mutability models for more details). We also used a mappability filter that describes whether sites can be readily aligned to the genome and removed both mutations and non-mutations at non-mappable sites as in previous work (Carlson et al. 2018).

### Genomic predictor data

To test which genetic and epigenetic properties of the genome predict mutability, we used publicly available tissue-specific data for numerous genomic predictors to train each of our mutability models. Each tissue-model contains the same predictor categories to facilitate comparison across tissues, including cell-type specific processes that may vary (e.g. histone modification, transcription) and genome sequence-based properties that are fixed across tissues (e.g. GC content). These genetic and epigenetic factors were selected based on prior evidence that they predict mutability (Michaelson et al. 2012; Chen et al. 2017; Carlson et al. 2018; García-Nieto et al. 2019).

It has been shown that some predictors affect mutation at different spatial scales in the genome.For example G/C bases tend to mutate at a higher rate than A/T but high GC at the scale of 1000bp tends to reduce spontaneous mutation (Ellegren et al. 2003; Supek and Lehner 2019). We therefore included the value of these properties across various spatial contexts. Predictors at multiple different scales were only included in the model where intercorrelation was low enough to permit (see supplemental tables S13-16 for VIF summaries). Supplemental table 2 contains a summary of the predictors used in our model.

Measurements of some tissue-specific predictors were vulnerable to technological inconsistencies that could bias our analysis. For example, transcription data measured with RNASeq across different tissues could have different depth of coverage, read length or read architecture. Thus we strictly standardised tissue specific predictors so that for a given predictor we chose datasets that were as similar as possible based on the molecular method, source lab, fragment length, read length, date submitted, file type, cell type, age, and sex. In-depth details regarding these predictors including the data source and lab technique can be found in table S2.

### Training mutability models

To explore which genomic or epigenetic properties drive mutation, we trained models that used these properties to predict whether a given site was a mutant or non-mutant in our training data. The relative probability (0 - 1.0) that a site would be in the mutation category is referred to as ‘mutability’. Extending previous work (Michaelson et al. 2012; Ness et al. 2015; Carlson et al. 2018), we used logistic regression to train a predictive model that used the genetic and epigenetic properties to distinguish between mutated and non-mutated sites using the following equation:

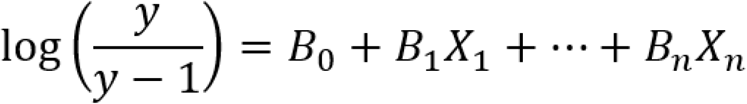

where y is the predicted mutability for a given site, B(0) represents the intercept, B(j) is the coefficient of predictor j, and X(j) is the value of predictor j in a given site. We checked model assumptions through visually assessing linearity (see figure S9-12) and tested for independence by calculating the variance inflation factor (VIF) for each predictor in each model (tables S4-S7).

Non-mutant sites were randomly selected from the mappable genome without overlap with the mutation dataset. There was an equal number of mutant and non-mutant sites in each tissue model which allows comparison across models because the expected probability that a site is a mutation or not in each tissue is 50%. The full dataset was split evenly into a training and testing set, with equal proportions of mutant and non-mutant sites in each half. Model accuracy was assessed on the independent “testing” dataset. All code used for these analyses is available at: https://github.com/MadeleineOman/mutability/tree/master

### Model predictions and performance

To assess how well our models predicted mutability, we used mean absolute error (MAE). MAE is generally calculated as the mean absolute difference between the predicted and actual response value of a given data point. In our case this would be whether a mutant site (1) is properly predicted to have a mutability >0.5 or a non-mutant site (0) with a mutability <0.5. However, mutation rate is a continuous property with some sites being more likely to mutate than others. Therefore we need to test whether sites that are predicted to have higher mutability are proportionally more likely to be mutations and vice versa. To do so we can aggregate all the sites in the test set with a given mutability value and determine if the fraction of mutations in that bin is proportional to mutability as follows:

1. Each site receives a predicted mutability value from the model.
2. Sites are ranked by predicted mutability.
3. Sites are grouped into bins based on their mutability (fixed 400 sites per bin).
4. Within each bin, the mean predicted mutability as well as the proportion of true mutations within the bin are calculated.
5. We plot the predicted vs mean observed mutability in each bin on a scatter plot (e.g. figure 1).
6. The fit of the line relating observed to predicted mutability is calculated

Using this method means that we may not specifically know the accuracy of any one site prediction nor how much variance there truly is in a bin (Eyre-Walker and Eyre-Walker 2014), but instead we can test if the mean mutability of the bin is accurate which allows us to interpret the fit and relative effect of predictors. Bin size was set at 400 sites per bin, as it is the maximum size that allows at least 20 bins in the smallest training dataset (blood n_sites_=9082). To test the validity of this binning method we compare accuracy to a null model where the model predicts on data where the mutation status of the sites are randomized so that there should be no genomic properties that predict mutability (see figure S1).

To explore how mutational processes vary between tissues, we also used trained models from each tissue to predict mutability of sites from other tissues. If a model presents high error when predicting on another tissue, this indicates differences in the strength and direction of predictors in the fitted model in each respective tissue.

### Genetic and epigenetic predictors of mutability

To understand which genomic properties and epigenetic processes are the best predictors of mutation we extracted the effect of each predictor from each tissue model. In regression, each predictor is assigned a β coefficient (slope) that describes the direction and strength of that predictor in determining mutability. The t-statistic of each predictor represents a normalized indicator of the importance of that predictor and more is suitable for comparison across models. The t-statistic for a given predictor is calculated as follows,

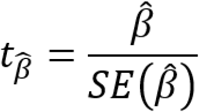

where β(hat) is the estimator for the true β value in the model and SE(β(hat)) is its standard error. To compare how the same genomic properties affected mutability differently across tissues, we normalized all predictors in all tissues to a mean of zero and standard deviation of one. This allowed us to directly compare the varying importance of each predictor in each tissue across models. T-statistic values for all predictors (as well as the beta coefficients, associated standard error,z-score, and p value) are available in supplementary table 9.

### Variation of tissue-specific mutability of individual sites

We also wanted to compare how much the mutability of any given site in the genome might vary depending on the tissue. To achieve this we chose 15000 random sites from the genome. For each site we used the trained tissue-specific mutability model and predictor data to estimate the site’s predicted mutability in each of the four tissues. We then compared these sites across tissues in a pairwise manner and analyzed the similarity of mutability with a linear model and reported the correlation coefficient (*R*^*2*^) as a measure of similarity.

### Validating the model on an independent dataset

It is possible that mutagenic signatures may arise due to technical artefacts, for example the method of single cell culture or sequencing technology. To test for the effect of such artefacts we assessed the accuracy of the blood model when predicting on an independently generated dataset of mutations in blood (n=9,082, see table S1 for details). We assume that if the true mutability of these tissues is the same, that differences in the model fit represent technical differences in the data generation or variation among samples.

## Data access

The full analysis is available online at GitHub at https://github.com/MadeleineOman/mutability and all data used in this study are from public databases, details of which can be found in supplemental table S1 and corresponding code in the GitHub repository.

## Acknowledgements

This work was supported by a Natural Sciences and Engineering Research Council (NSERC) Discovery grant (RGPIN/06331-2016) and Canadian Foundation for Innovation John R. Evans Leaders fund (35591) to R.W.N. and a Canada Graduate Scholarship-Doctoral (CGS D) to M.O.

