## Supplementary material for "Comparing the predictors of mutability among healthy human tissues inferred from mutations in single cell genome data": Supplmental figures S1-S9 and tables S3-S8

### Supplemental Information

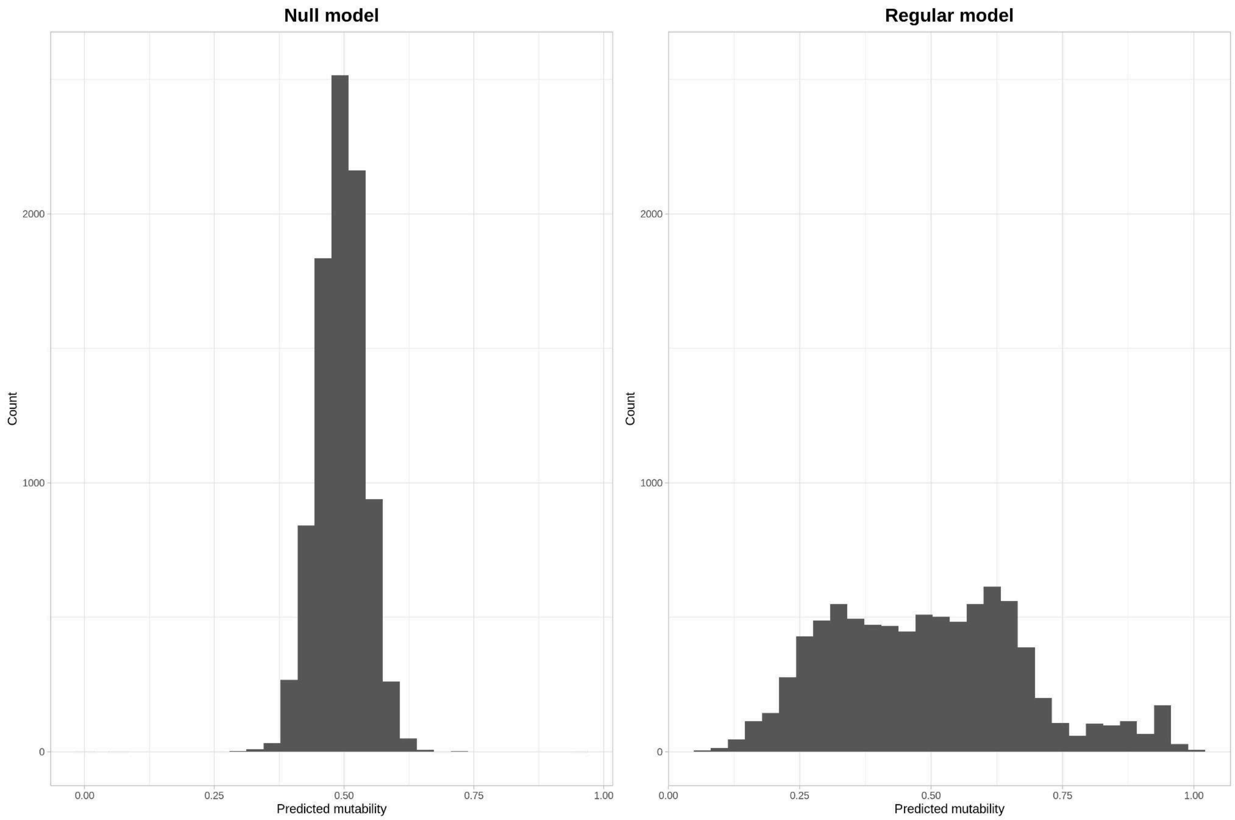


Figure S1: Comparison of model predictions in a null (left) and regular (right) model. Plotted is a histogram of the predictions made from the blood tissue model. The null blood model is identical as the regular blood model, except that it is trained on a dataset where mutations are randomized.

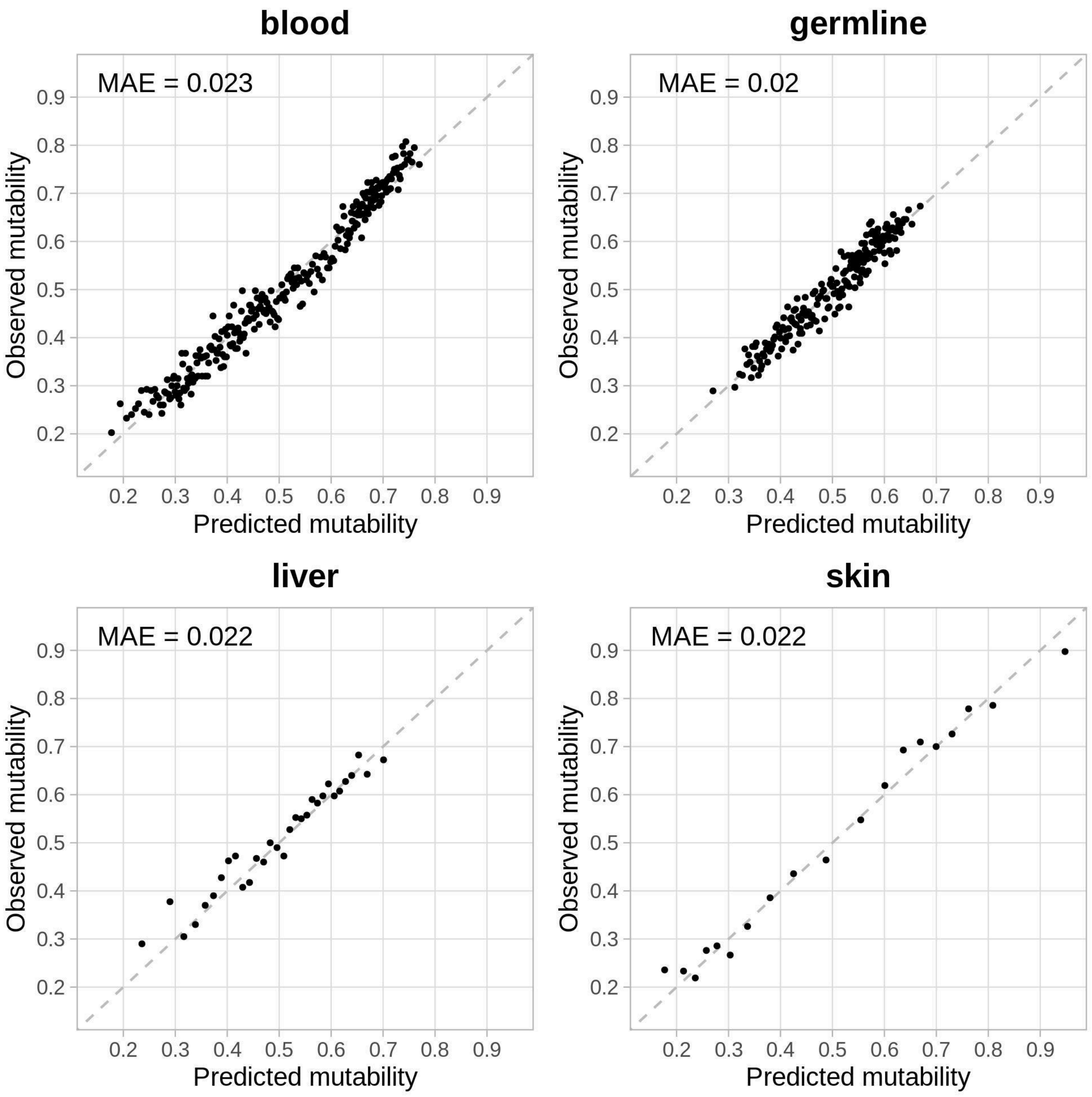


Figure S2: Tissue specific mutability model fit and accuracy in healthy tissues when CpG sites are removed from analyses. Here accuracy is estimated from how well the estimated mutability predicts the proportion of the sites in a given bin that are actually mutations. Each data point represents a bin of 400 mutated or unmutated sites within a given mutability range (n=400). The proportion of sites that are mutations in that bin is plotted against predicted mutability, which represents the model's estimation of the probability that a site would be a mutation in the test set, averaged for all sites in the bin (error bars = standard error of mutability in a bin). The dashed grey line is the one-to-one line, where the predicted mutability and proportion of mutations are equal.

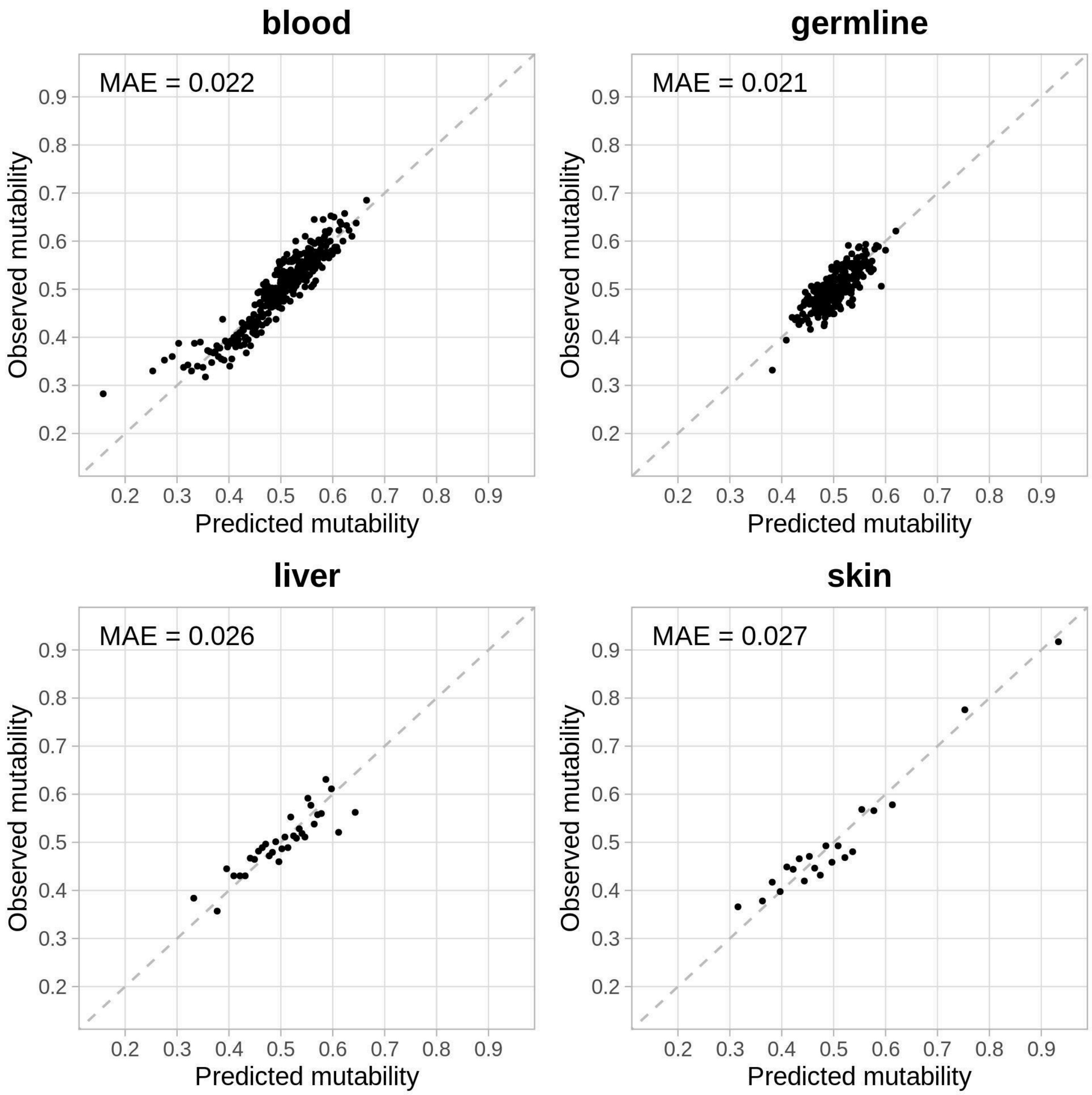


Figure S3: Tissue specific mutability model fit and accuracy in healthy tissues when triplets are not included as predictors. Here accuracy is estimated from how well the estimated mutability predicts the proportion of the sites in a given bin that are actually mutations. Each data point represents a bin of 400 mutated or unmutated sites within a given mutability range (n=400). The proportion of sites that are mutations in that bin is plotted against predicted mutability, which represents the model's estimation of the probability that a site would be a mutation in the test set, averaged for all sites in the bin (error bars = standard error of mutability in a bin). The dashed grey line is the one-to-one line, where the predicted mutability and proportion of mutations are equal.

Table S3: Comparing the spread and MAE of bins between the full model, and the model without triplet predictors.

| | Percent decrease in spread of predictions | Percent increase in MAE |
|---|---|---|
| blood | 50% | 10% |
| germline | 46% | 16% |
| liver | 53% | -10% |
| skin | 24% | 14% |
| Mean | 43% | 7.5% |

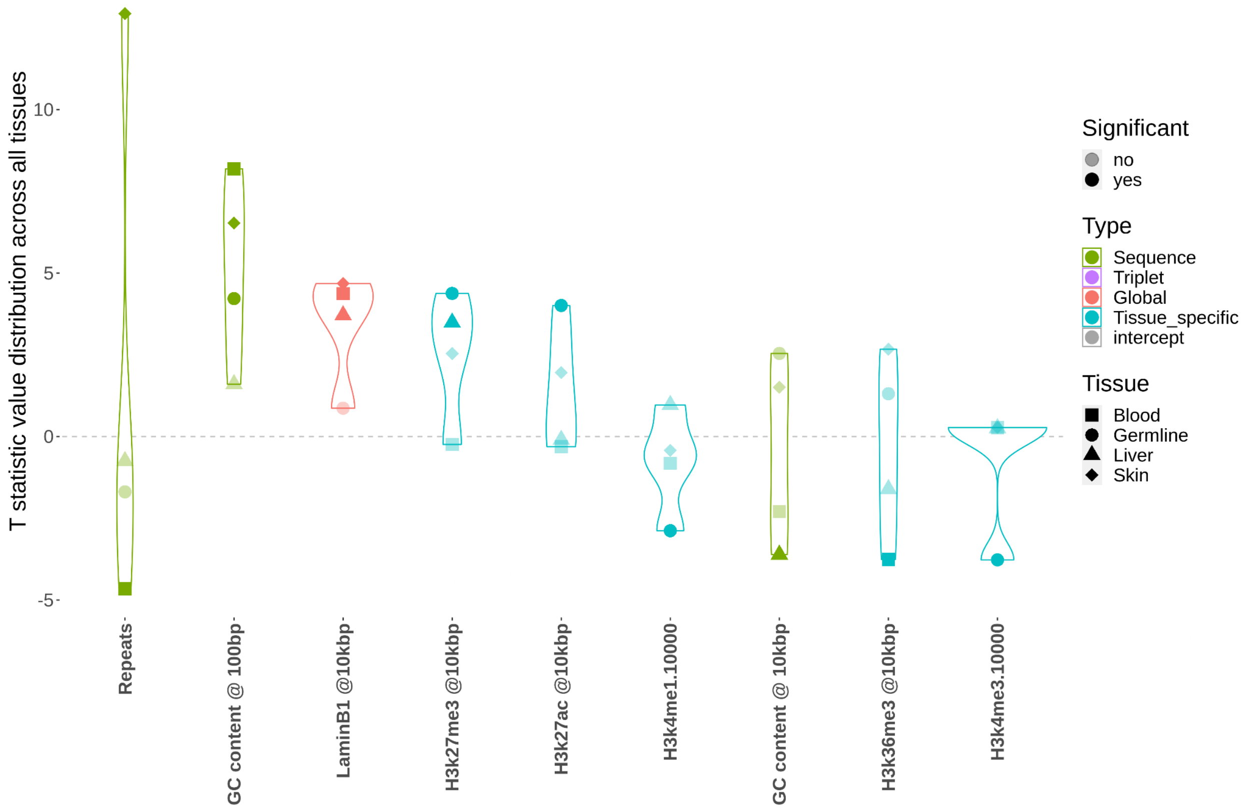


Figure S4: Comparison of the importance of genomic and epigenetic predictors of mutability among tissues in models without triplet predictors. Every predictor is represented by four points corresponding to each of the tissues, where each point represents the t-statistic for a given tissue. The larger deviations from zero (grey dotted line) indicate a stronger effect on mutability, with points above the line increasing mutability and those below reduce mutability. Here we only plot predictors that have significant effect ($p < 0.005$) in at least one tissue; tissues in which a given genomic property has no significant effect are indicated with translucent data points. Predictors are ordered by the most extreme (ie largest absolute value) t-statistic value of the four tissues. A predictor with the same mutagenic importance in two tissues would exhibit a larger t-statistic in the tissue with the larger dataset. Thus, to ensure discrepancies in t-statistics reflect biologically meaningful differences, here the germline, liver and skin datasets are downsampled to be equivalent to the smallest dataset, skin ($n_{mutations}$ = 10,191).

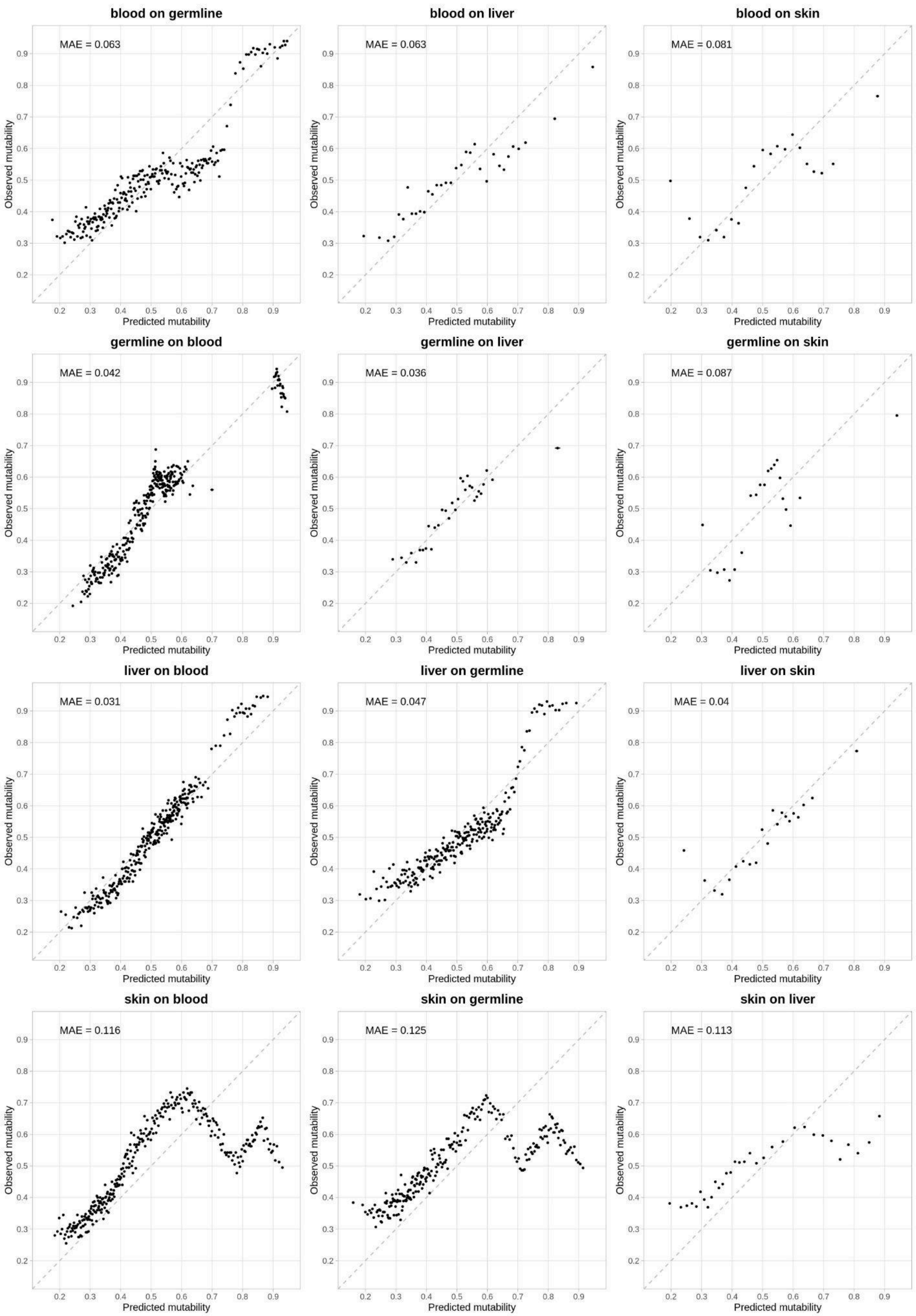

blood on germline
MAE = 0.063
blood on liver
MAE = 0.063
blood on skin
MAE = 0.081
germline on blood
MAE = 0.042
germline on liver
MAE = 0.036
germline on skin
MAE = 0.087
liver on blood
MAE = 0.031
liver on germline
MAE = 0.047
liver on skin
MAE = 0.04
skin on blood
MAE = 0.116
skin on germline
MAE = 0.125
skin on liver
MAE = 0.113
Observed mutability
Predicted mutability

Figure S5: Tissue specific mutability model fit and accuracy in healthy tissues, where the model predicts mutations from tissues other than the one on which the model was trained. Here accuracy is estimated from how well the estimated mutability predicts the proportion of the sites in a given bin that are actually mutations. Each data point represents a bin of 400 mutated or unmutated sites within a given mutability range (n=400). The proportion of sites that are mutations in that bin is plotted against predicted mutability, which represents the model's estimation of the probability that a site would be a mutation in the test set, averaged for all sites in the bin (error bars = standard error of mutability in a bin). The dashed grey line is the one-to-one line, where the predicted mutability and proportion of mutations are equal.

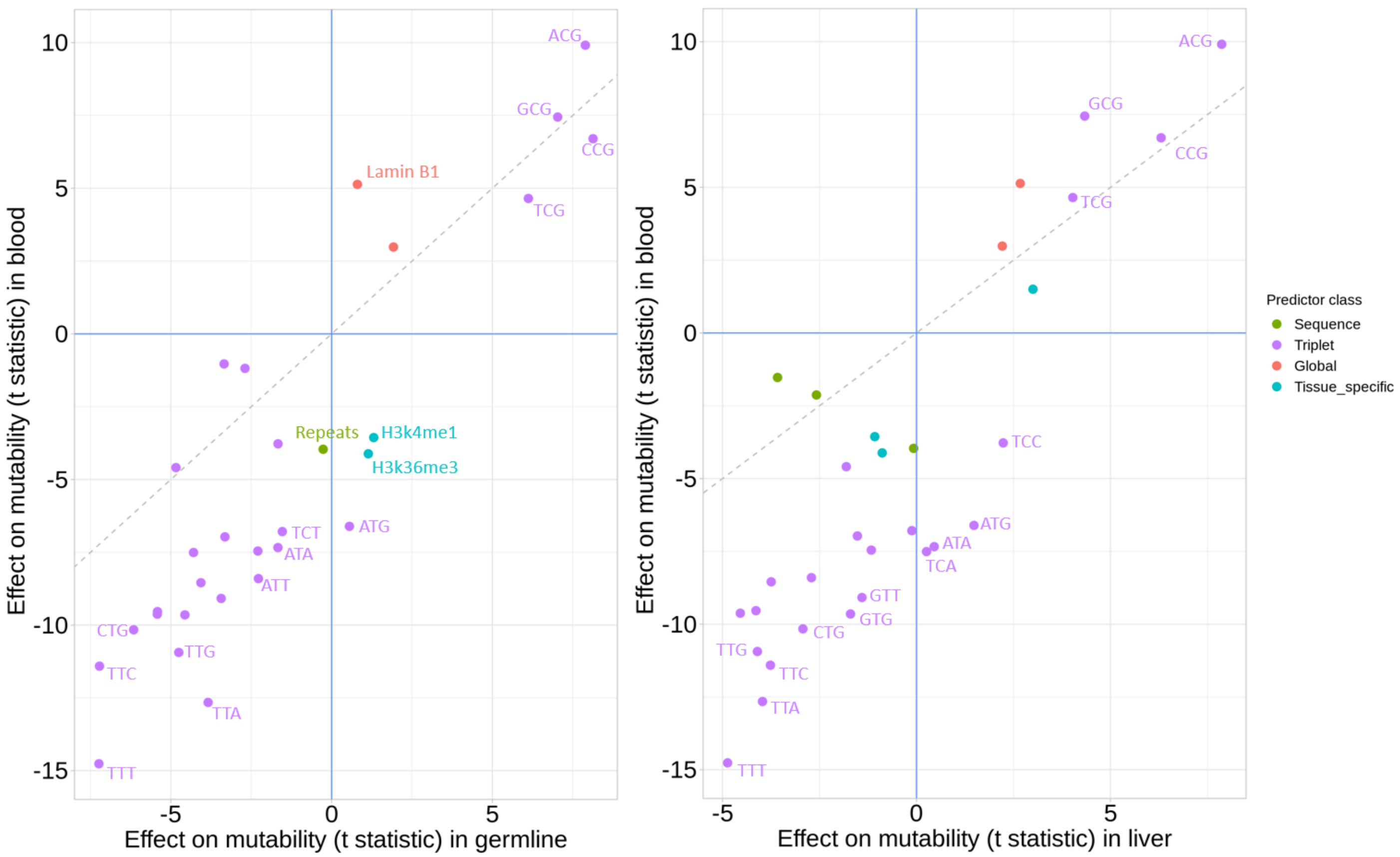


Figure S6: Pairwise comparisons of the strength and direction of different genetic and epigenetic properties and processes on mutability between blood and germline (A), and blood and liver (B). Only predictors that are significant in at least one of the tissues ($p<0.01$) are displayed. Predictors that have a similar effect in both tissues are near 1:1 line (grey dotted line), while those that deviate have an effect unique to that tissue. Predictors in the top left or bottom right quadrant change the direction of effect between tissues, possibly indicating a process that is mutagenic in one tissue and protective in another. Select informative points are labelled. A predictor with the same mutagenic importance in two tissues would exhibit a larger t-statistic in the tissue with the larger dataset. Thus, to ensure discrepancies in t-statistics reflect biologically meaningful differences, here the germline, liver and skin datasets are downsampled to be equivalent to the smallest dataset, blood (n mutations = 9,082).

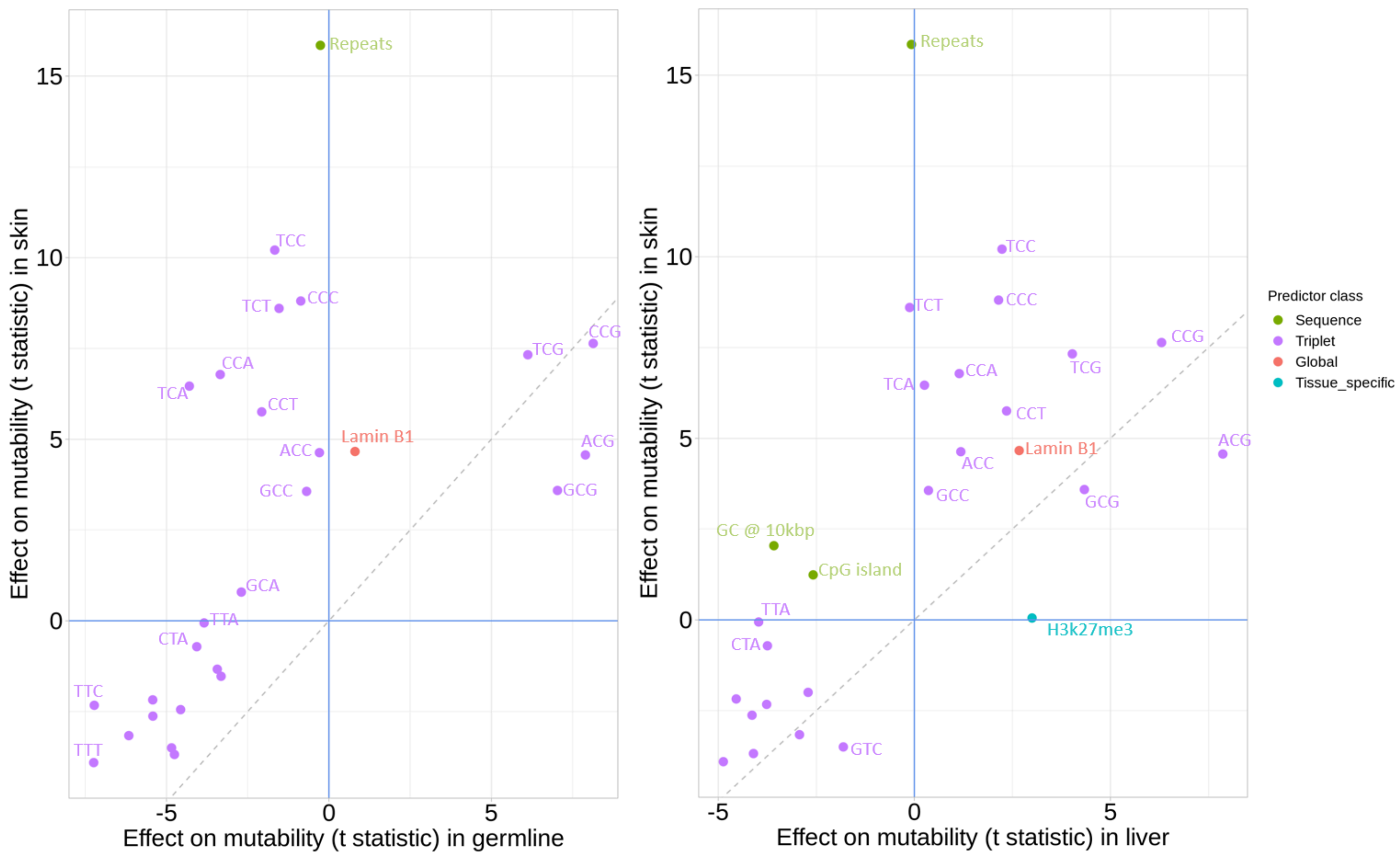


Figure S7: Pairwise comparisons of the strength and direction of different genetic and epigenetic properties and processes on mutability between skin and germline (A), and skin and liver (B). Only predictors that are significant in at least one of the tissues ($p<0.01$) are displayed. Predictors that have a similar effect in both tissues are near 1:1 line (grey dotted line), while those that deviate have an effect unique to that tissue. Predictors in the top left or bottom right quadrant change the direction of effect between tissues, possibly indicating a process that is mutagenic in one tissue and protective in another. Select informative points are labelled. A predictor with the same mutagenic importance in two tissues would exhibit a larger t-statistic in the tissue with the larger dataset. Thus, to ensure discrepancies in t-statistics reflect biologically meaningful differences, here the germline, liver and skin datasets are downsampled to be equivalent to the smallest dataset, blood (n mutations = 9,082).

Table S4: Blood VIF values that exceed 2.

| Predictor names | Adjusted R squared | VIFs |
|---|---|---|
| H3k27.10000 | 0.787231 | 4.69993 |
| H3k4me1.10000 | 0.702406 | 3.360285 |
| GC_content.10000 | 0.597471 | 2.484291 |
| H3k4me3.10000 | 0.549842 | 2.221444 |

Table S5: Germline VIF values that exceed 2.

| Predictor names | Adjusted R squared | VIFs |
| --- | --- | --- |
| H3k27.10000 | 0.747547 | 3.961137 |
| GC_content.10000 | 0.636493 | 2.750978 |
| H3k4me1.10000 | 0.633515 | 2.728627 |
| H3k4me3.10000 | 0.617543 | 2.614672 |
| tripletTTT | 0.524473 | 2.10293 |

Table S6: Liver VIF values that exceed 2.

| Predictor names | Adjusted R squared | VIFs |
| --- | --- | --- |
| H3k27.10000 | 0.702681 | 3.363388 |
| H3k4me3.10000 | 0.58663 | 2.419143 |
| GC_content.10000 | 0.577271 | 2.365582 |
| tripletTTT | 0.533105 | 2.141808 |
| tripletTCT | 0.502957 | 2.011898 |

Table S7: Skin VIFs values that exceed 2.

| Predictor names | Adjusted R squared | VIFs |
|---|---|---|
| GC_content.10000 | 0.663297 | 2.969973 |
| tripletTCT | 0.639599 | 2.774691 |
| tripletTCC | 0.630876 | 2.709119 |
| H3k4me1.10000 | 0.59423 | 2.464449 |
| tripletCCT | 0.567217 | 2.310625 |
| tripletTCA | 0.560977 | 2.277784 |
| tripletTTT | 0.56023 | 2.273915 |
| tripletCCC | 0.548682 | 2.215733 |
| tripletCCA | 0.54858 | 2.215233 |
| GC_content.100 | 0.537397 | 2.161683 |

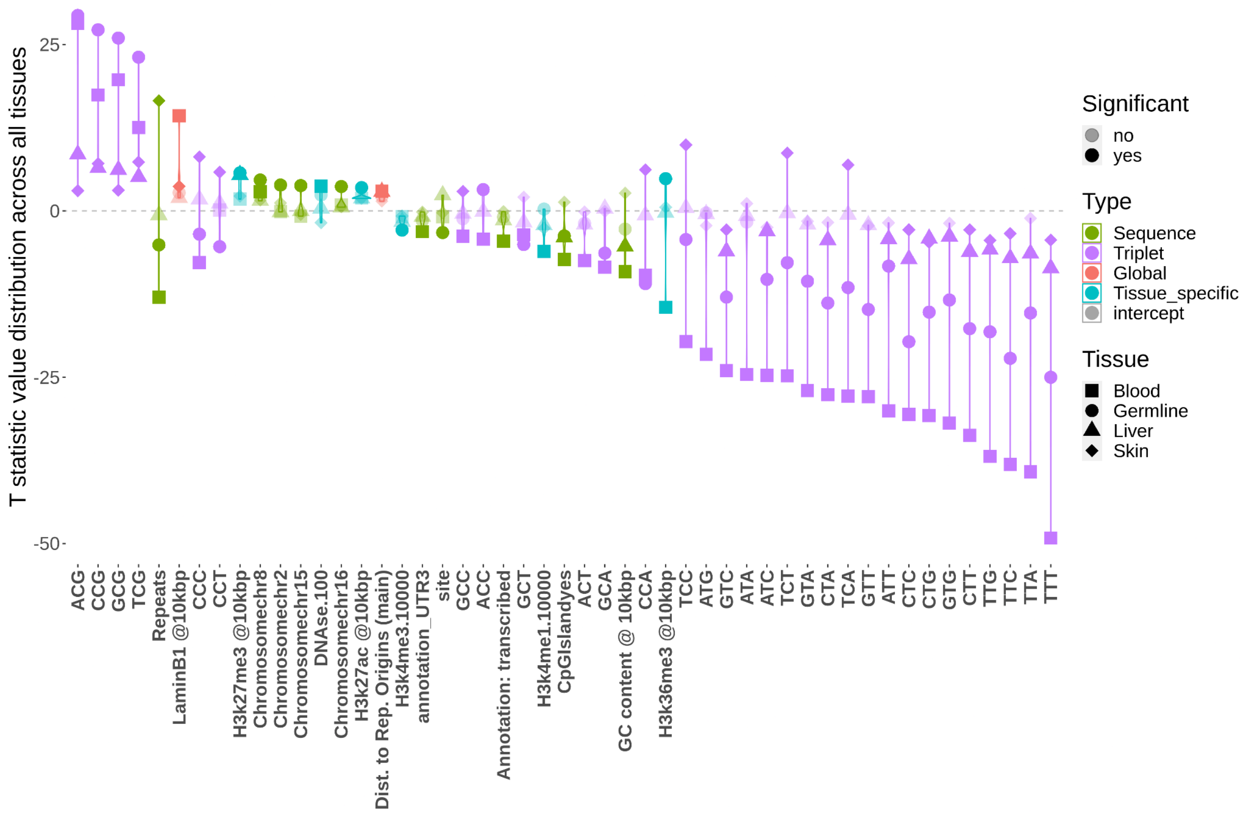


Figure S8: Comparison of the importance of genomic and epigenetic predictors of mutability among tissues in models that have not been downsampled. Every predictor is represented by four points corresponding to each of the tissues, where each point represents the t-statistic for a given tissue. The larger deviations from zero (grey dotted line) indicate a stronger effect on mutability, with points above the line increasing mutability and those below reduce mutability. Here we only plot predictors that have significant effect (p <0.005) in at least one tissue; tissues in which a given genomic property has no significant effect are indicated with translucent datapoints. Predictors are ordered by the most extreme (ie largest absolute value) t-statistic value of the four tissues.

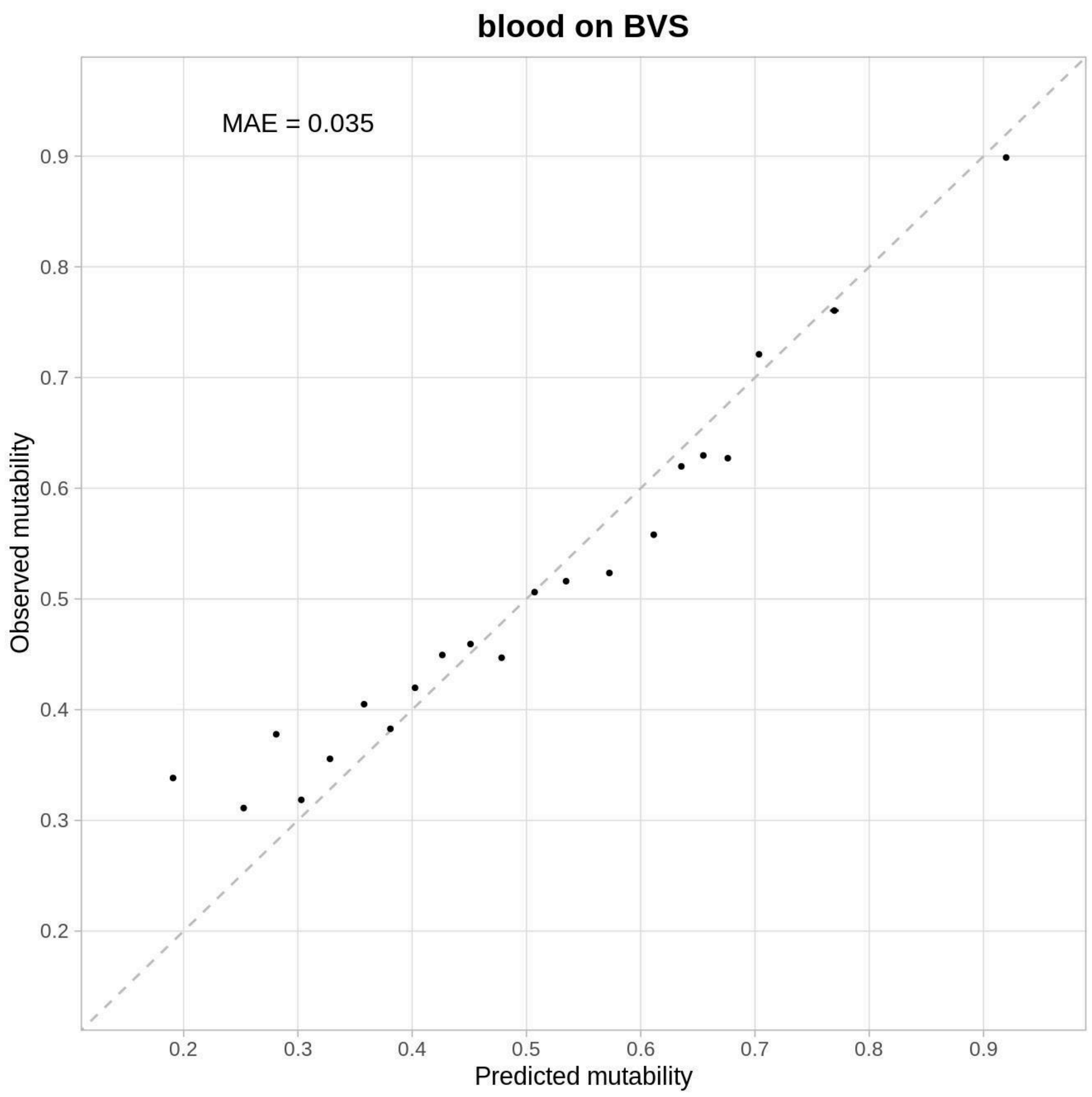


Figure S9: Validation of the blood model on the blood validation set (BVS): an independently generated dataset of single cell mutations in blood. Tissue specific mutability model fit and accuracy in healthy tissues, where the model predicts mutations from tissues other than the one on which the model was trained. Each data point represents a bin of sites which includes both mutated and unmutated sites (n=400). The proportion of sites that are mutations in that bin is plotted against predicted mutability, which represents the model's estimation of the probability that a site would be a mutation in the test set, averaged for all sites in the bin. The dashed grey line is the one-to-one line, where the predicted mutability and proportion of mutations are equal. Full details on the BVS dataset are available in supplemental table 1.

Table S8: Details of which labels sourced from the annotation tracks (see table S2 for full details) are sorted into the categories included in our model “transcribed”,”UTR3”,and ”UTR5”. Sites that do not have a label are categorized as “untranscribed” in our model.

| Model label | Included labels |
|---|---|
| “UTR3” | “three_prime_UTR” |
| “UTR5” | “five_prime_UTR” |
| “transcribed” | "antisense_RNA", "CDS", "exon", "gene", "lnc_RNA", "mRNA", "ncRNA", "miRNA", "tRNA", "vault_RNA","V_gene_segment", "Y_RNA", "RNase_MRP_RNA", "RNase_P_RNA", "rRNA", "scRNA", "snoRNA", "snRNA",”telomerase_RNA", "transcript", "primary_transcript", "cDNA_match", "CDS", "exon","gene", "transcript", "start_codon", "stop_codon", "D_gene_segment" , stop_codon_redefined_as_selenocysteine", "J_gene_segment", "C_gene_segment" |
